# Structural Mechanics of the Alpha-2-Macroglobulin Transformation

**DOI:** 10.1101/2021.09.22.461418

**Authors:** Yasuhiro Arimura, Hironori Funabiki

## Abstract

Alpha-2-Macroglobulin (A2M) is the critical pan-protease inhibitor of the innate immune system. When proteases cleave the A2M bait region, global structural transformation of the A2M tetramer is triggered to entrap the protease. The structural basis behind the cleavage-induced transformation and the protease entrapment remains unclear. Here, we report cryo-EM structures of native- and intermediate-forms of the *Xenopus laevis* egg A2M homolog (A2Moo or ovomacroglobulin) tetramer at 3.7-4.1 Å and 6.4 Å resolution, respectively. In the native A2Moo tetramer, two pairs of dimers arrange into a cross-like configuration with four 60 Å-wide bait-exposing grooves. Each bait in the native form threads into an aperture formed by three macroglobulin domains (MG2, MG3, MG6). The bait is released from the narrowed aperture in the induced protomer of the intermediate form. We propose that the intact bait region works as a “latch-lock” to block futile A2M transformation until its protease-mediated cleavage.

## Introduction

Alpha-2-Macroglobulin (A2M) family proteins are multi-functional proteins that are highly conserved among metazoans and are also found in archaea and bacteria (Budd et al., 2004; Garcia-Ferrer et al., 2017; Zorin and Zorina, 2017). While a variety of A2M functions are known in diverse biological processes, including those as a regulator of cytokines and growth factors and those as a protein chaperon to bind to misfolded proteins such as amyloid beta proteins associated with Alzheimer’s disease (Cater et al., 2019), the conserved function of A2M family proteins is protease inhibition, which contributes to the innate immune system via counteracting pathogenic proteases (Wong and Dessen, 2014). Metazoan genomes encode multiple A2M homologs/paralogs such as pregnancy zone protein (PZP), α1-inhibitor-3 (α1I3), the complement components C3, C4, and C5, the cell surface antigen CD109, and the complement C3 and PZP-like α2M domain-containing 8 (CPAMD8). These A2M family proteins belong to the thioester-containing proteins (TEPs), sharing a common evolutionary origin and conserved structural and functional features (Garcia-Ferrer et al., 2017). TEPs are commonly comprised of more than seven macroglobulin-like domains (MG), C1r/C1s, Uegf, and Bmp1 found domain (CUB), and thioester domain (TED). Despite their conserved domain architecture, multiple oligomerization states (monomer, dimer, tetramer, or octamer) of A2M family proteins are reported depending on the subtypes (Garcia-Ferrer et al., 2017; Zorin and Zorina, 2017).

Since Barrett and Starkey proposed almost 50 years ago that cleavage of a sensitive peptide of A2M by the prey protease irreversibly transforms the A2M structure and traps the prey (Barrett and Starkey, 1973), two distinct protease trapping mechanisms have been proposed; the “Venus flytrap” mechanism that only works in tetrameric A2M family proteins (Marrero et al., 2012) (Figure S1A), and the “snap trap” mechanism that can work with a monomeric form (Garcia-Ferrer et al., 2015) (Figure S1B).

In the Venus flytrap mechanism, proteases are physically trapped inside the hollow “prey chamber” of the A2M tetramer, which is abundantly found in serum and egg white (Ikai et al., 1983; Marrero et al., 2012; Miller and Feeney, 1966; Zorin and Zorina, 2017) (Figure S1A). While many versions of the Venus flytrap models have been proposed (Feldman et al., 1985; Harwood et al., 2021; Kolodziej et al., 1996, 2002; Larquet et al., 1994; Marrero et al., 2012; Sottrup-Jensen, 1989; Tapon-bretaudiere et al., 1985), it is commonly assumed that a native-form A2M tetramer undergoes a global transformation when a protease cleaves the bait region within the BRD (bait-region domain), entrapping the protease inside the closed chamber of the induced form. However, the structural basis behind the Venus flytrap mechanism remains highly speculative since the 3D structure of the native-form A2M tetramer has never been solved. During 1970-80s, it was reported that the native-form A2M compacts upon binding to proteases, based on native PAGE, small-angle X-ray scattering (SAXS), and negative-stain electron microscopy (EM) (Barrett and Starkey, 1973; Barrett et al., 1979; Österberg and Pap, 1983; SCHRAMM and SCHRAMM, 1982; Tapon-bretaudiere et al., 1985). From these earlier studies, Feldman et al predicted that the native-form A2M tetramer forms a hollow cylinder structure with two open ends (Feldman et al., 1985), whereas Sottrup-Jensen proposed that the A2M tetramer forms a cross-like structure where two rodlike dimers are bridged by disulfide-bonds (Sottrup-Jensen, 1989). A more recent computational modeling predicted that the native-form A2M tetramer forms a hollow cylinder-like structure with two ~40 Å open ends where a protease can enter the prey chamber (Marrero et al., 2012). However, earlier biochemical analyses showed that larger proteases such as plasmin (approximately 90 x 60 x 50 Å) could be captured by A2M (Pizzo and Roche, 1987). This cannot be explained by the hollow cylinder-like structure with the 40 Å entry sites. In the latest atomic model of the native A2M tetramer constructed by negative-stain EM, SAXS, and cross-linking–mass spectrometry (XL-MS) (Harwood et al., 2021), the native-form A2M tetramer was predicted to form a hollow tube-like structure, where each protomer would twist roughly a half-turn upon transformation into the induced-form tetramer (Harwood et al., 2021). However, it is not clear how the bait cleavage induces these twisting motions of each molecule. To answer this fifty-year-old question about the Venus flytrap mechanism, high-resolution structure information of the native-form A2M tetramer is needed.

Meanwhile, the monomeric A2M homologs, such as bacterial A2M and mosquito TEP1r can also capture and inactivate proteases (Garcia-Ferrer et al., 2015, 2017; Le et al., 2012; Wong and Dessen, 2014), indicating that A2M family proteins have another protease inhibition mechanism distinct from the Venus flytrap mechanism. This “snap trap” mechanism is also triggered by the bait region cleavage but traps a protease by forming a covalent β-cysteinyl γ-glutamyl thioester bond with the protease at the conserved CXEQ (X=G or L) motif in the TED domain (Wong and Dessen, 2014) (Figure S1B). In the absence of substrates, the reactive thioester on the CXEQ motif is protected by surrounding aromatic and hydrophobic residues on the TED and RBD domains by forming a hydrophobic pocket that is conserved in bacterial and eukaryotic A2M family proteins (Janssen et al., 2005; Wong and Dessen, 2014). The reactive thioester is released from the hydrophobic pocket upon the global A2M structural transformation caused by the proteolytic cleavage of the flexible bait region, which is recognized by a variety of endo-proteases (Wong and Dessen, 2014).

For both the “Venus trap” and “snap trap” mechanisms, the key questions are how the native form tetramer is prevented from a futile, spontaneous structural transformation, and how the bait region cleavage irreversibly triggers the structural transformation of the A2M tetramer. To answer these questions, native- and induced-form A2M tetramer structures, especially around the bait region, must be compared. However, with X-ray crystallography, fair structural comparisons of the native- and induced-form A2Ms under similar conditions is technically challenging since the global structural transformation of A2Ms does not allow the same crystal packing (Goulas et al., 2014). In addition, due to its flexible nature, the bait region structure has never been observed in the previously reported X-ray crystallography studies of native-form *Salmonella enterica ser*. *Typhimurium* A2M monomer (Wong and Dessen, 2014), native-form mosquito TEP1r monomer (Le et al., 2012), induced-form *E. coli* A2M monomer (Garcia-Ferrer et al., 2015), and induced-form human A2M tetramer (Marrero et al., 2012). In the native-form human C3 monomer, which harbors the anaphylatoxin (ANA) and α’NT domains in place of BRD, the structures of these domains are partially determined (Janssen et al., 2005), and the conformational change of α’NT upon the cleavage of the ANA has been reported (Janssen et al., 2005, 2006). Nevertheless, it is not clear if this mechanism is conserved in the bait region/BRD of the A2M tetramer.

We recently reported a 5.5 Å-resolution cryo-EM structure of the protein complex that we assigned as a tetrameric form of A2M family proteins co-fractionated with the nucleosomes assembled in *Xenopus* egg extract (Arimura et al., 2021). Since the structure was distinct from the crystal structure of induced-form human A2M tetramer, we assumed that they represent the native-form tetramers. In this study, by comprehensively re-analyzing the same cryo-EM dataset, we determined structures of the native-form and intermediate-form tetramers at 4.1 Å and 6.4 Å resolutions, respectively. Further local refinement on the protomer in the native-form tetramer improves the resolution to 3.7 Å. By “reading” amino acid side chain structures, we identify the protein as “Uncharacterized protein LOC431886 isoform X1” and rename it A2Moo or ovomacroglobulin. While 2D structures of the native-form A2Moo tetramer structure are consistent with most previous EM studies, the 3D structure does not match to previously proposed hollow cylinder models. Moreover, we successfully determine the location of the bait region in the native-form tetramer, leading us to propose a latch-lock model, which can explain how the global transformation of A2M is triggered by bait region cleavage.

## Result

### Cryo-EM structure determination of A2Moo, the A2M family protein in *Xenopus* egg extract

In our recent study, by applying the Template-, Reference- and Selection-Free (TRSF) cryo-EM pipeline for nucleosome-enriched fractions isolated from the interphase and metaphase chromosomes formed in *Xenopus* egg extracts, we fortuitously co-determined the 5.5 Å resolution cryo-EM structure of tetrameric A2M family proteins (Arimura et al., 2021). The structure did not match to any reported structures, including the crystal structure of the induced-form human A2M tetramer, which is more compacted than our structure (Marrero et al., 2012). Because it has been established that the native-form A2M tetramer exhibits more open structure than the induced form (Barrett et al., 1979; Tapon-bretaudiere et al., 1985), and because our A2M was isolated from the functional egg cytoplasm, where ovochymase (the major egg serine protease secreted upon fertilization) is inactive (Lindsay et al., 1999, 1992), we assumed that this A2M structure represents the native form tetramer. In this study, we aimed to obtain higher resolution structures of these A2M tetramers and their structural variants (Figure S2). To pick more A2M family protein particles from the existing micrographs (EMPIAR-10691, EMPIAR-10692, EMPIAR-10746, and EMPIAR-10747), we employed the machine learning-based particle picking software Topaz (Bepler et al., 2019). In addition to the apparent native-form A2M tetramer, the 3D classification isolated an intermediate-form tetramer in which one out of four A2M protomers is structurally different from the native form (Figure 1A). Structures of the native- and intermediate-form tetramers were determined at 4.1 Å and 6.4 Å resolutions, respectively ((Figure 1A, S2, and S3A). Further local refinement on the protomer revealed the 3.7 Å resolution structure ((Figure 1A, S2, and S3A). In the 3.7 Å resolution cryo-EM map, the amino acid side chains are distinguishable, allowing us to deduce the amino acid sequence identified from the cryo-EM map (Figure 1B). Amongst sixteen A2M family proteins encoded in the *Xenopus laevis* genome, “Uncharacterized protein LOC431886 isoform X1” (GenBank accession; XP_018079932) satisfied the cryo-EM map (Figure 1B, Table S1). Gene ID for LOC431886 is ovos2.L, whose name infers ovostatin. However, the amino acid length and sequence of LOC431886 is closer to *Xenopus laevis* alpha-2-macroglobulin (XP_018080172.2) than to *Xenopus laevis* ovostatin (XP_041426100.1) (Figure S3A, Table S2). Similarly, LOC431886 is closer to human alpha-2-macroglobulin than to human ovostatin homolog 1 or ovostatin homolog 2 (Figure S3A, Table S2). Moreover, LOC431886 possesses the CXEQ motif, which is missing in ovostatin homologs in chickens and humans (Fig. S3C). As the Xenbase proteomics analysis reports that LOC431886 is dominantly abundant in oocytes and eggs during embryonic development (Peshkin et al., 2019), we renamed LOC431886 protein A2Moo or ovomacroglobulin to stand for the major A2M variant in oocytes/eggs. Using the sequence information of A2Moo, atomic models were built for both native- and intermediate-forms of A2Moo.

**Figure 1.**
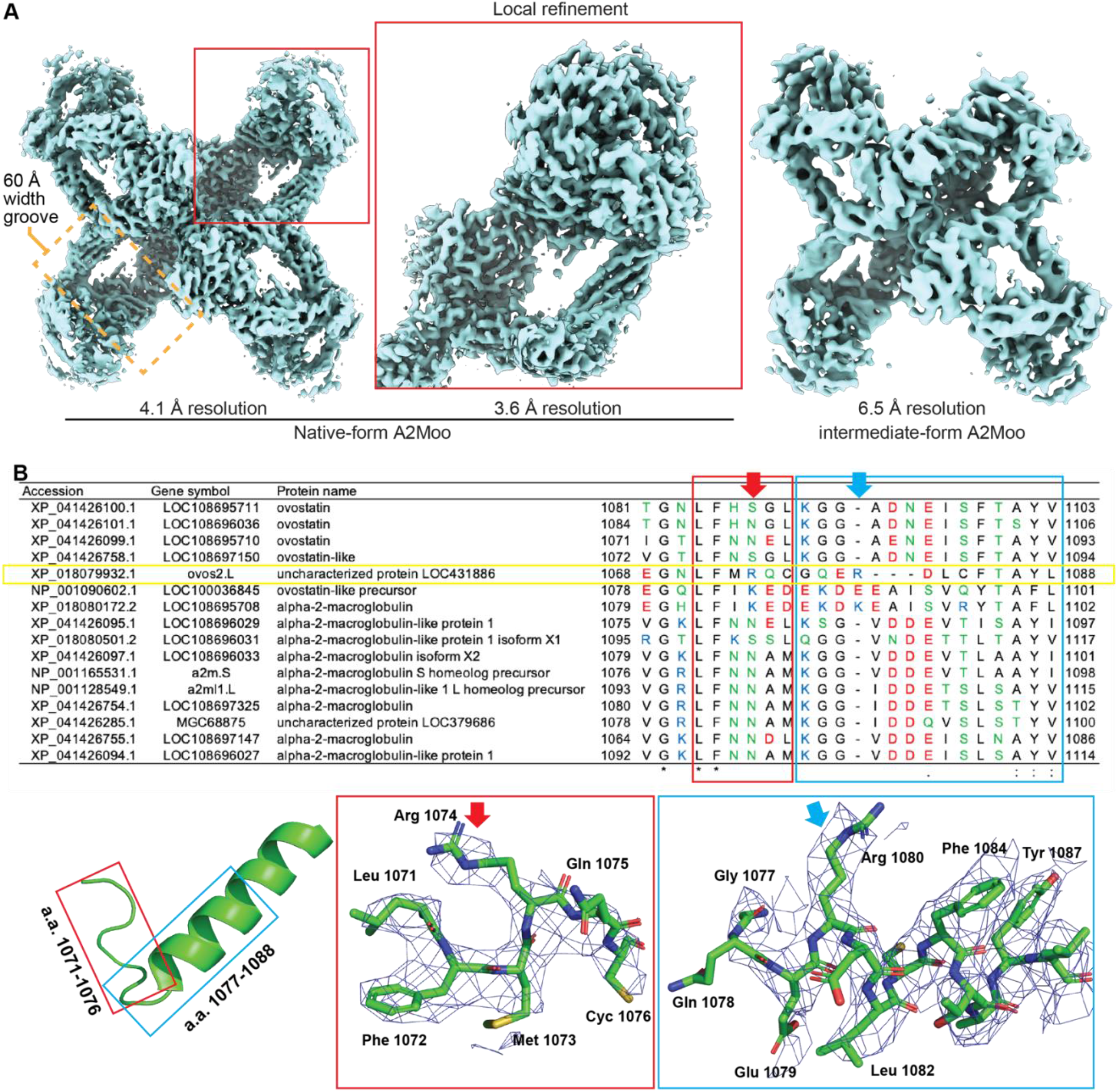
Cryo-EM structure determination of A2Moo. (**A**) Cryo-EM maps of the native-form A2M family protein tetramer (left), locally refined native-form A2M family protein protomer (center), and intermediate-form A2M family protein tetramer (right). (**B**) Identification of A2Moo, the A2M family protein in *Xenopus* egg. Top panel; amino acid sequence alignment of sixteen *Xenopus laevis* A2M family proteins with reasonable protein length to satisfy the EM map. The representative region used for protein identification is shown. A yellow rectangle indicates the protein that matches to the EM density (LOC431886: named A2Moo). Bottom panels; overlay of the atomic model of A2Moo and cryo-EM density of the locally refined native-form A2M protein protomer.

### The structural transformation of A2Moo tetramer

The 2D averaged class structures of native-form A2Moo tetramer mimics previously reported low-resolution 2D electron micrograph images of native-form ovomacroglobulin from crocodile egg white (Ikai et al., 1983), and those of the native-form A2M tetramers from mammalian sera (depicted as “Ж”, “four-petaled flower”, “eye”, “lip”, and “padlock”) (Bretaudiere et al., 1988; Larquet et al., 1994; Tapon-bretaudiere et al., 1985) (Figure 2A and S4A).

**Figure 2.**
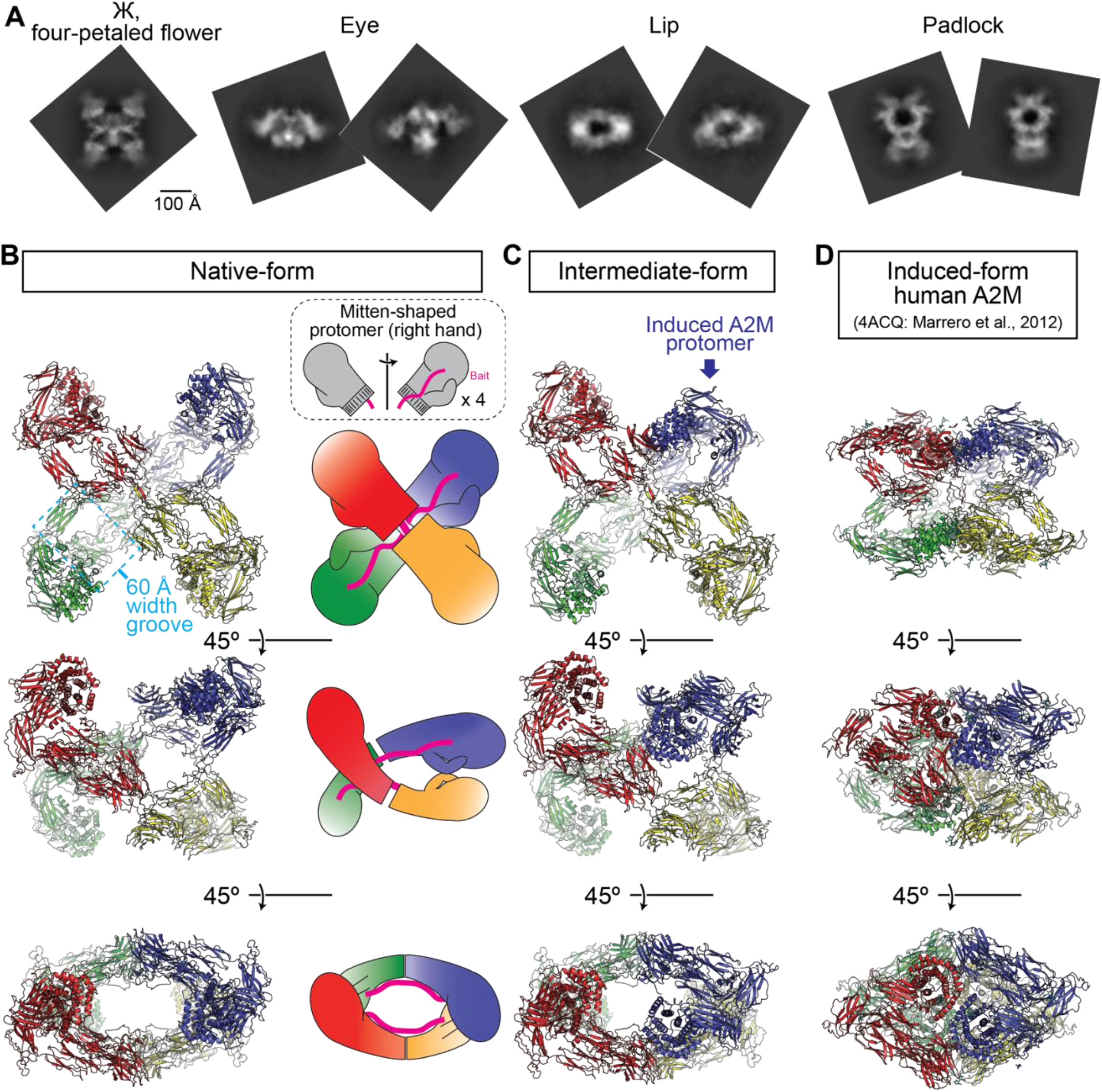
2D and 3D structures of the A2Moo tetramers. (**A**) Representative 2D class averages of the native-form A2Moo tetramer mimic previously proposed depictions of A2M architectures. (**B**) 3D atomic model of the native-form A2Moo tetramer. Two pairs of “connected mitten”-shaped A2M dimers stack to form a cross-like configuration with D2 symmetry, where each monomer consists of a “bulky finger” module, a “thumb” module, and a “palm” module. (**C**) 3D atomic model of the intermediate-form A2Moo tetramer. The “bulky finger” module folds toward the “wrist”. (**D**) Atomic model of the induced-form human A2M (PDB ID: 4ACQ) (Marrero et al., 2012).

In the 3D structure of the native-form A2Moo tetramer, two pairs of “connected mitten”-shaped A2M dimers stack to form a cross-like configuration with D2 symmetry, where we define three structural modules for each monomer; a “bulky finger” module, a “thumb” module, and a “palm” module (Figure 2B, Movie S1). Each monomer is linked to another protomer at the opposite end through their “wrists” (chains A-B, and chains C-D), as well as linked laterally to the second protomer through its thumb (chains A-C, and chains B-D). Interestingly, among the numbers of 3D models, to the best of our knowledge, only a hand-written model by Sottrup-Jensen in 1989 and a low-resolution 3D map reconstructed by Larquet et al in 1994 captured this overall shape (Figure 2B) (Larquet et al., 1994; Sottrup-Jensen, 1989), but not by recent computationally simulated atomic models of A2M native forms (Harwood et al., 2021; Marrero et al., 2012). In our native structure, four large 60 Å-wide grooves, each containing an exposed bait region (see Figure 3B, Movie S1), can be identified. For example, a 60 Å-wide groove is surrounded by the bulky finger domain and the thumb domain of Chain A, the thumb domain of Chain C, and the palm domain of the Chain D. Adjacent pairs of these grooves are connected to form two large chambers (Movie S1).

**Figure 3.**
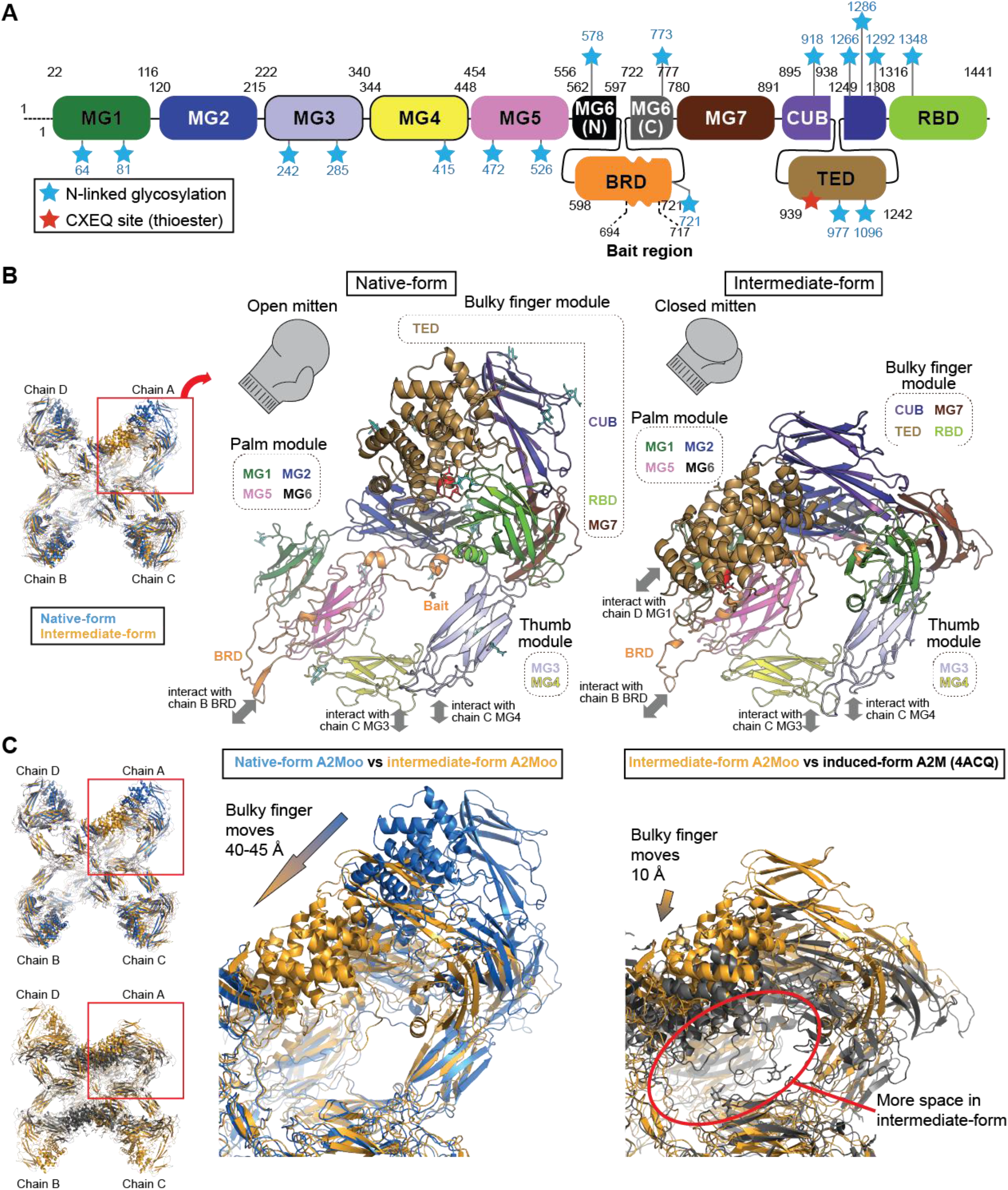
3D arrangements of the A2Moo domains. (**A**) Domain organization of A2Moo. (**B**) 3D arrangements of the domains of the native-form A2Moo (left) and an induced-protomer of the intermediate-form A2Moo (right). (**C**) Structural comparison between a protomer of the native-form and induced-protomer of the intermediate-form A2Moo (left) and between the induced-protomer of the intermediate-form frog A2Moo and a protomer of the induced-form human A2M (PDB: 4ACQ) (right).

Each native-form A2Moo monomer consists of eleven structural domains (MG1-MG7, BRD, CUB, TED, and RBD) like the crystal structure of the induced-form human A2M (Marrero et al., 2012) (Figure 3A and 3B). The bulky finger module consisting of (TED+CUB+RBD+MG7) is attached to the palm module (MG1+MG2+MG5+MG6) and the thumb module (MG3+MG4), while a stretched BRD (in chain A) links its bulky finger module to another BRD (in chain B) at the opposite end (or the “wrist”). The palm module does not laterally interact with another protomer, whereas the thumb module (in chain A) dimerizes with another thumb module (in chain C). In the native-form A2Moo, a reactive β-cysteinyl-γ-glutamyl thioester on the CXEQ motif in the TED domain is protected by surrounding hydrophobic- and aromatic-residues on TED and RBD (Figure S4B). In addition, seventeen *N*-linked glycosylation on asparagine residues at Asn64, Asn81, Asn242, Asn285, Asn415, Asn472, Asn526, Asn578, Asn721, Asn773, Asn918, Asn977, Asn1096, Asn1266, Asn1286, Asn1292, and Asn1348 are observed (Figure S5).

In the intermediate-form, the structure of one of four A2Moo protomers (chain A) is dramatically rearranged. The structure of this “induced” A2Moo protomer is similar to the crystal structure of induced-form human A2M (Fig. 2C and 2D) (Marrero et al., 2012), while structural changes on the other protomers are minor. The intermediate-form A2Moo tetramer structure implies that the structural transformation within one A2M protomer does not necessarily trigger the global structural transformation of the tetramer. While the induced protomer also consists of eleven domains, the 3D arrangement of these domains is different from the native-form protomer (Figure 3B and 3C, Movie S1). Most prominently, in the transformed form, the bulky finger module folds toward the wrist, resulting in ~45 Å slide of CUB and TED. This large movement is accompanied by the dissociation of TED from RBD, and the generation of a new lateral interaction between TED and MG1 of the adjacent chain D protomer, closing the groove (Figure 3 and S6, Movie S2). This lateral TED-MG1 (or interprotomeric bulky finger-palm) interaction is also observed in the induced-form human A2M crystal structure (Marrero et al., 2012), which was transformed by methylamine (but see Discussion). The bulky finger modules (MG7+CUB+TED+ RBD) of induced-form human A2M further slide ~10 Å toward the center of the tetramer compared to the induced-form protomer in the intermediate A2Moo tetramer (Figure 3C right). In the induced-form protomer, the CXEQ site is exposed to solvent, facing inside the prey chamber (Figure 3B).

### The path of the bait region in the native-form A2Moo

It was not clear how A2M structural transformation is induced upon bait cleavage since the structure of the bait region in the native form A2M tetramer has never been solved. Even in the crystal structures of native-form monomeric A2M homologs, the flexible nature of the bait region made the electron density ambiguous (Le et al., 2012; Wong and Dessen, 2014). The cryo-EM density of the flexible part of the bait region (a.a. 694-717) was also ambiguous in the locally refined 3.7Å resolution cryo-EM map of the native-form A2Moo (Figure 4A and 4B). However, with cryo-EM single particle analysis, the structure of the flexible region is often visualized in a low-pass filtered map, even though it is averaged out in the high-resolution map. Indeed, the path of the bait region can be traced in the 11 Å-resolution low-pass filtered map of the native-form A2Moo (Figure 4C). The bait region of A2Moo is located beside the MG5 and MG6 along with the N terminal part of BRD in the palm module (Figure 4C and 4D).

**Figure 4.**
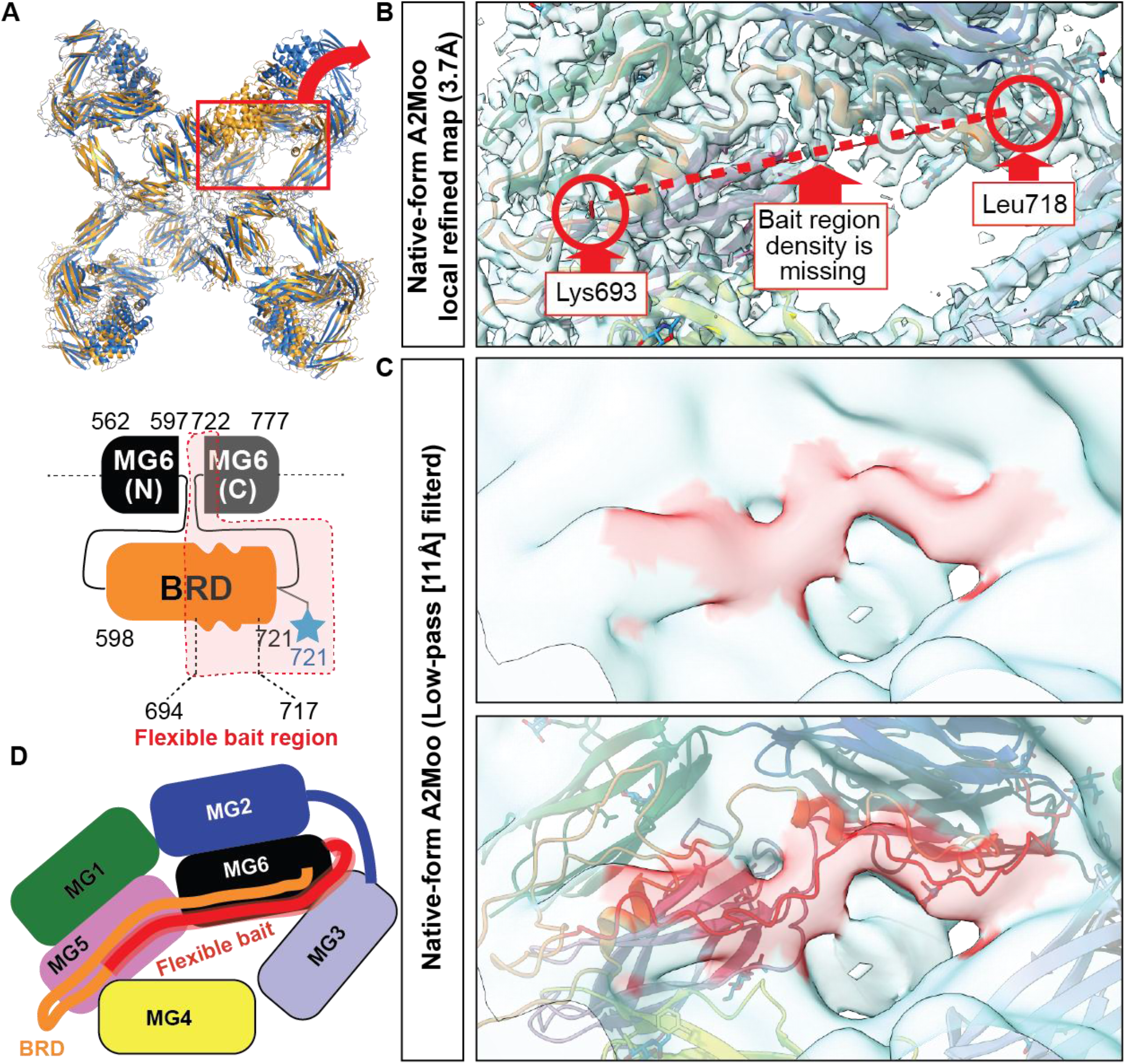
The path of the bait region in the native-form A2Moo. (**A**) Tetramer atomic model and cartoon model to depict the location of flexible bait region. (**B**) High-resolution locally refined map around flexible bait region. The density of the bait region is missing. (**C**) Low-pass (11Å) filtered map around flexible bait region. The bait region of A2Moo can be traced. (**D**) The cartoon representation depicting the location of the flexible bait region. The bait region of A2Moo is located beside the MG5 and MG6.

### Structural transformation around the “latch hole”

In the native-form A2Moo, the C-terminal part of the bait domain threads through an aperture composed of MG2, MG3, and MG6 (Figure 5A and 5B). We call this aperture a “latch hole” for the following reasons. EM density of the C-terminal side of the bait region (a.a. 718-726, red), which threads through the latch hole, is observed in the 3.7 Å locally refined native-form A2Moo map (Figure 5B). However, EM density corresponding to the bait region is not observed within the latch hole in the induced protomer of the intermediate-form A2Moo (Figure 5C). Moreover, in the induced protomer, MG2 and MG6 are shifted to fill the latch hole (Figure 5C). These data suggest that the bait region is cleaved in the induced protomer and comes out from the latch hole (Figure 5D). A similar structural transformation was also observed in the complement C3 crystal structures, in which the bait region is replaced with ANA and α’NT regions (Janssen et al., 2005, 2006); the α’NT region of C3 comes out from the small aperture between MG2, MG3, and MG6 in C3b (induced-form with ANA removal) or C3c (induced-form with ANA and TED removal), suggesting that this structural transformation mechanism is conserved between A2M and C3. Since bait region cleavage is responsible for the A2M transformation, we hypothesize that threading of the intact bait region into the latch hole locks the movement of MG2 and MG6, but protease-mediated bait cleavage unlocks the MG2 and MG6 and consequently induces the observed global structural transformation of A2M (Figure 5D, Movie S1). Intriguingly, since the reactive thioester-containing TED domain is located next to MG2, the shifted MG2 and MG6 domains upon induction may push TED away from RBD, which protects thioester on TED by hydrophobic interaction in the native form (Figure 5D, Movie S1).

**Figure 5.**
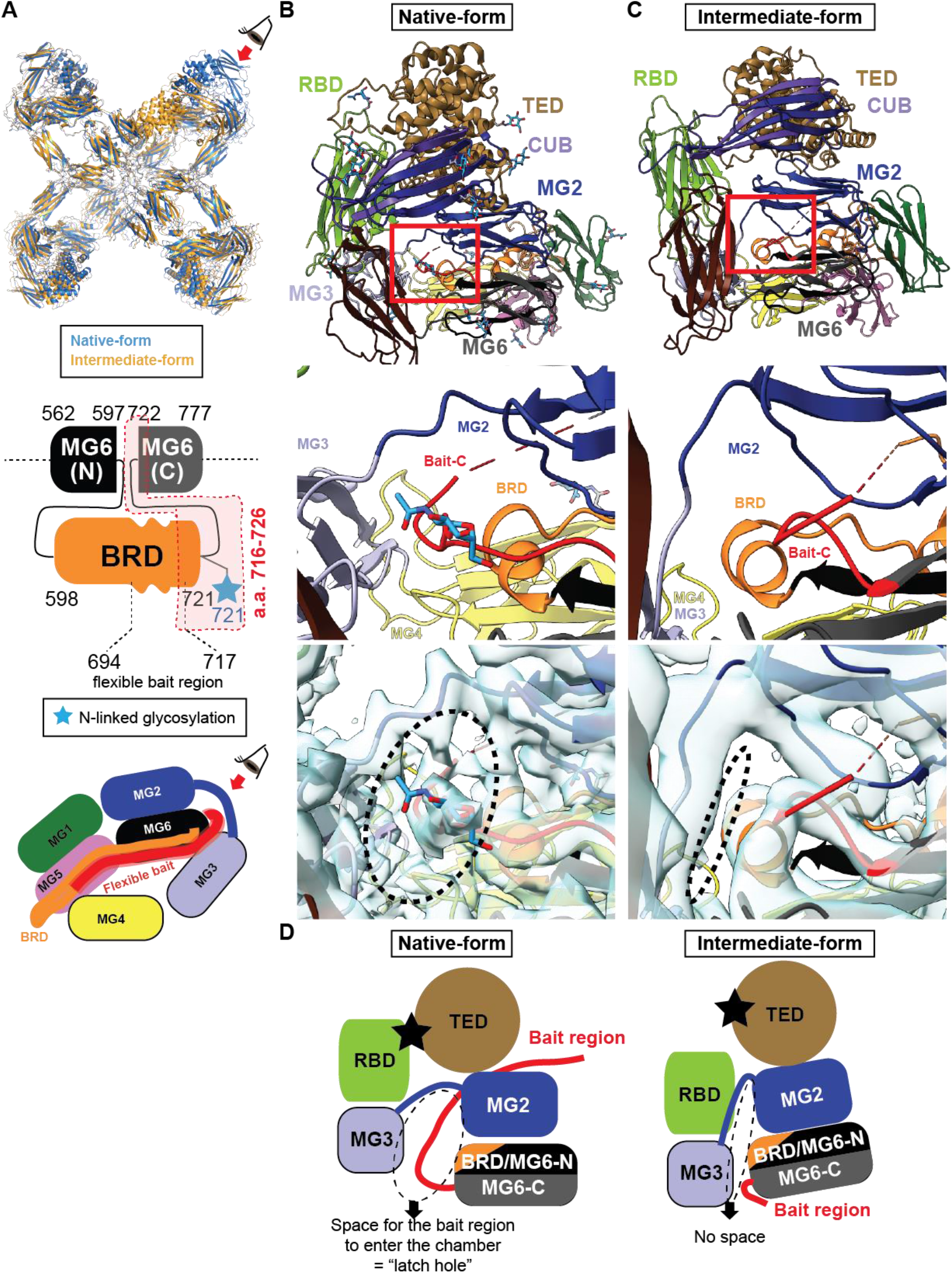
Structural transformation around the “latch hole”. (**A**) Tetramer atomic model and cartoon model to depict the viewpoint in Figure 5. (**B, C**) Structures around the latch hole in a protomer of the native-form A2Moo (B) and induced-protomer of the intermediate-form A2Moo (C). Top panels show a protomer in A2Moo tetramers. Middle panels show a zoom-up view around the latch hole. (**D**) The cartoon representation depicting the structural transformation around the latch hole. In the induced protomer, the bait region is not observed within the latch hole, and MG2 and MG6 are shifted to fill the latch hole.

### Structural variations of native-form A2Moo tetramer

In the cryo-EM map of the native-form A2Moo tetramer, EM density around the bulky finger module (MG7, CUB, TED, and RBD) is ambiguous as compared to other regions (Figure 1A), suggesting their structural flexibility. To capture structural variations of these regions, we performed 3D variability analysis (3DVA) in CryoSPARC (Punjani and Fleet, 2021), which captures structural variations in the cryo-EM data as trajectories of motion. Each trajectory represents a principal component of the major structural variations. In Figure 6A and Movie S2, four types of structural variations are shown. In each structural variation, the bulky finger module flexibly moves in many directions (Figure 6A and Movie S3). In addition, the framework of the prey chambers also flexibly moves in many ways (Figure 6A and Movie S2). These flexible movements may help the protease accessibility to the bait region. For example, in the “open” native-form A2Moo structure, 60Å diameter sphere and model structure of plasmin (90 kDa) can easily access the bait region (Figure 6B and 6C).

**Figure 6.**
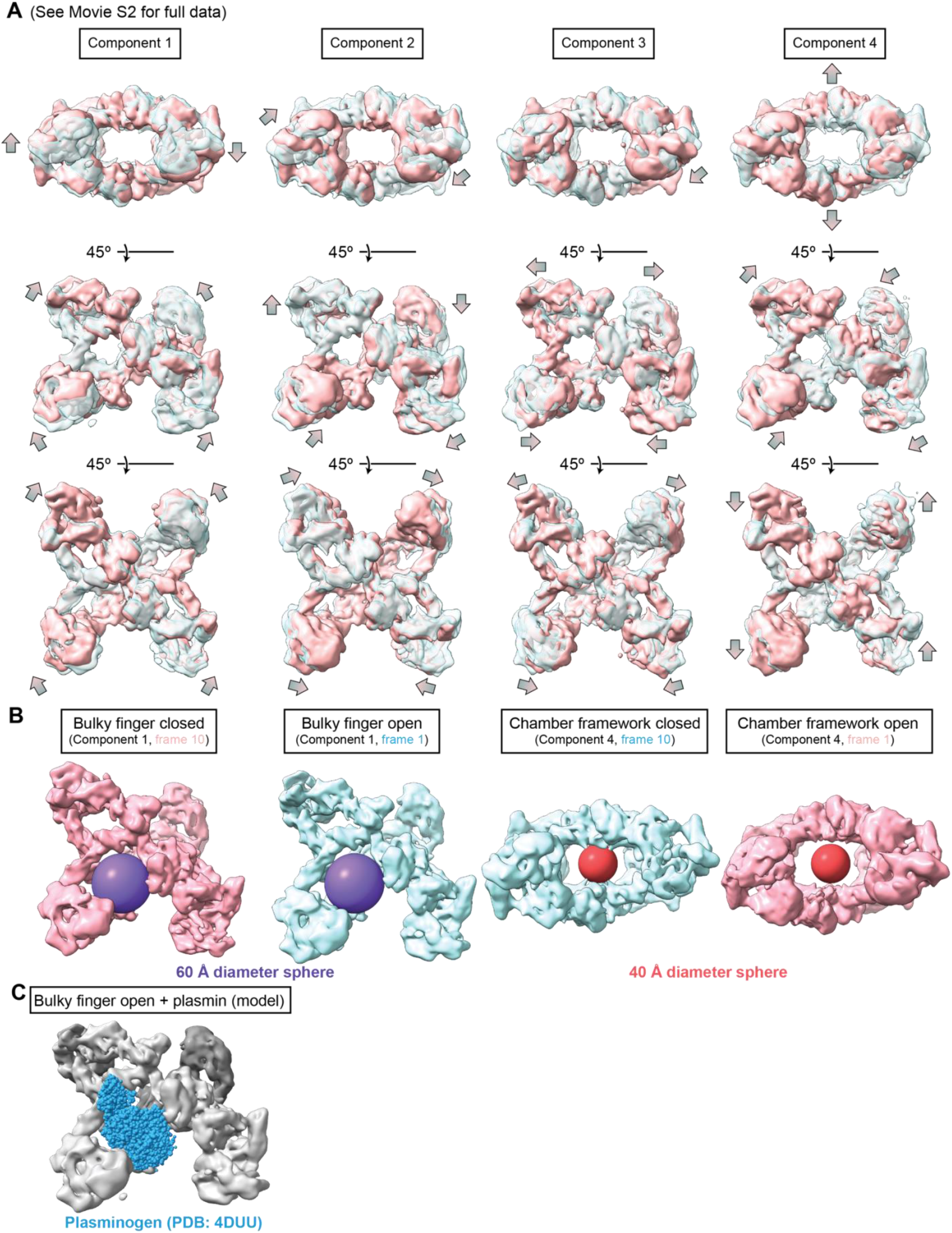
Structural variations of native-form A2Moo tetramer. (**A**) 3DVA of the native-form A2Moo tetramer. Four components of the significant structural variations are shown. The bulky finger modules flexibly move in many ways. (**B**) The flexible A2M variation expands the 60 Å groove and prey chamber. (**C**) Structural model of the plasmin serine protease domain accessing the bait region of the 60 Å groove in native-form A2M tetramer. Instead of plasmin, full length plasmin precursor (human plasminogen) structure (PDB: 4DUU) was mapped on the “open” native-form A2Moo tetramer (Law et al., 2012).

## Discussion

### Native-form A2M tetramer structure

In this study, we solved high-resolution cryo-EM structures of the native- and intermediate-form A2Moo tetramer from *Xenopus* eggs. The overall 3D structure of the native-form A2Moo tetramer is distinct from the previously proposed hollow cylinder models (Fig. S1A) (Feldman et al., 1985, Kolodziej et al., 2002, Marrero et al., 2012, Harwood et al., 2021a), but is consistent with the coarse models depicted by Sottrup-Jensen and Larquet et al more than 25 years ago (Fig. S1B)(Larquet et al., 1994; Sottrup-Jensen, 1989), and recaptures a variety of 2D views of the native A2M reported in old electron micrographs (Bretaudiere et al., 1988; Ikai et al., 1983; Larquet et al., 1994; Tapon-bretaudiere et al., 1985). Our cryo-EM analyses show that the native-form A2M tetramer has two large protease chambers, each of which is composed of two connected 60 Å grooves formed between a pair of facing, unlinked protomers. The flexible nature of A2M tetramer may allow the access of proteases even larger than the 60 Å sphere (Figure 6 and 7A). Furthermore, whereas both Harwood’s and Marrero’s simulations predicted that all four protomers of the A2M tetramers are transformed simultaneously (Harwood et al., 2021; Marrero et al., 2012), the existence of the intermediate structure indicates that structural conversion of one protomer may not be necessarily sufficient to trigger the global transformation (Figure 7A).

**Figure 7.**
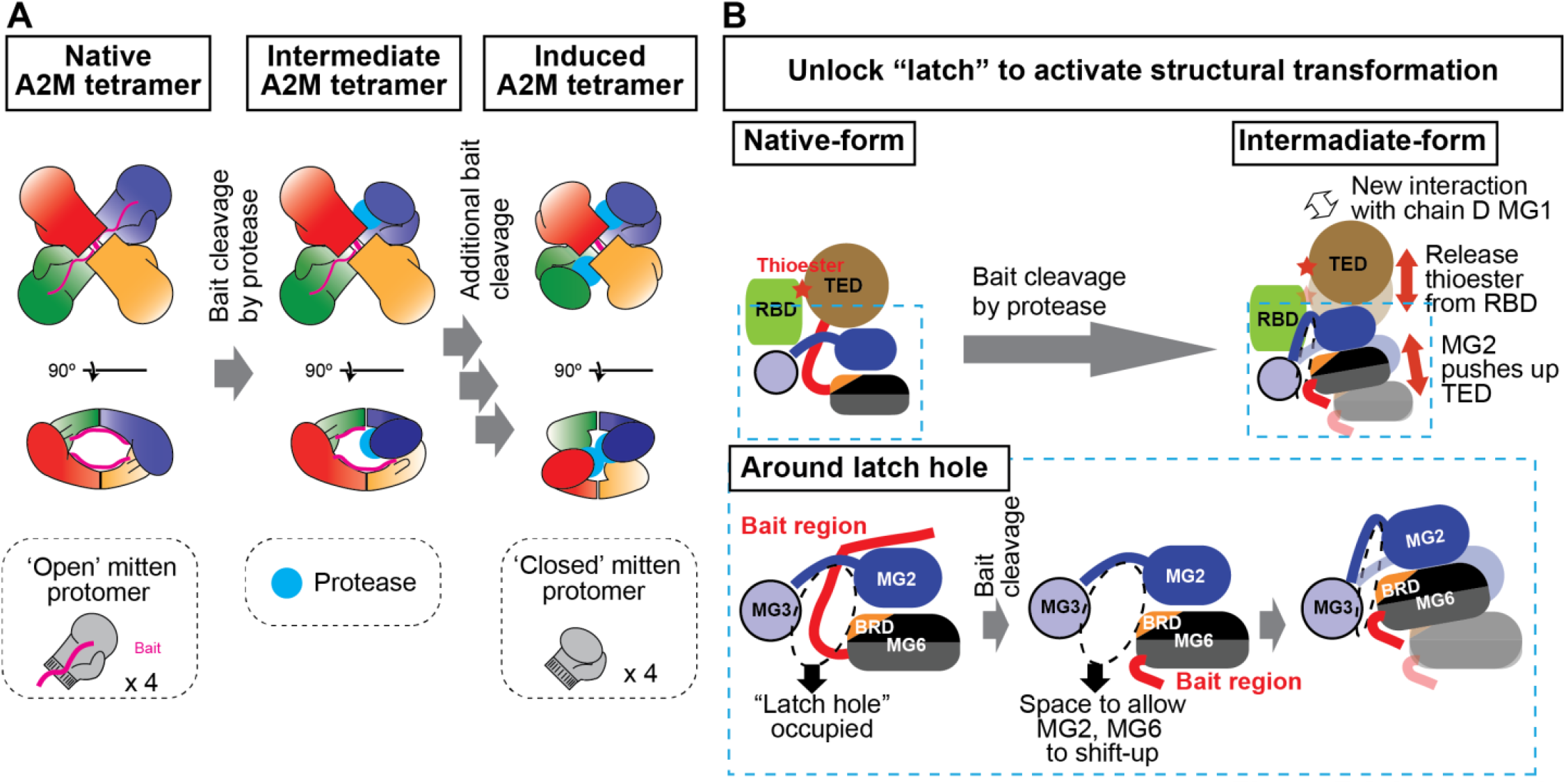
The model mechanics of the A2M transformation during protease inactivation. (**A**) Protease entrapment by A2M tetramer by the Venus flytrap mechanism. The cross-like structure and flexible nature of native-form A2M tetramer allow large proteases to access the bait region of A2M inside the 60 Å groove. In the intermediate-form tetramer, an A2M protomer is induced transformation by bait cleavage. In the induced-form tetramer, proteases can be trapped in the prey chamber. (**B**) Mechanics of A2M tetramer structural transformation by bait cleavage. The intact bait region in the latch hole blocks the shifting of MG2 and MG6, while bait cleavage unlocks this movement. The shifting of MG2 and MG6 pushes TED away from the RBD. The released TED makes a new interaction with MG1 of the adjacent native-form protomer to stabilize the induced-form A2M.

### Mechanics of A2M tetramer structural transformation by bait cleavage

For both the “Venus flytrap” and “snap trap” mechanisms, the key question was how the bait region cleavage induces the structural transformation of A2M. In this study, identification of the bait region path in native-form A2Moo provides us with a clue (Figure 4 and 5). The bait region is located beside the MG5 and MG6 in the palm module along with the N terminal part of the BRD and linked to the MG6 domain by threading through the aperture (or latch hole) formed with MG2, MG5, and MG6 (Figure 4C and 5B), in a way similar to the path of the α’NT region in monomeric human C3 (Janssen et al., 2005). In the induced-protomer in the intermediate-form A2M, the bait region does not exist in the latch hole, which is narrowed due to the sliding of MG2 and MG6 (Figure 5C). This suggests that the intact bait region in the latch hole blocks the shifting of MG2 and MG6, while bait cleavage unlocks the movement (Figure 7B). The shifting of MG2 and MG6 may also push TED away from the RBD that protects the reactive thioester on TED by hydrophobic interaction (Figure 7B). Although the path of the bait region is different from our model, Garcia-Ferrer et al proposed a similar model by comparing between an induced-form *E. coli* A2M (ECAM) monomer crystal structure and a low-resolution cryo-EM structure of native-form ECAM (Garcia-Ferrer et al., 2015). Therefore, this snap trap mechanism may be conserved among monomeric and tetrameric A2M family proteins.

In addition, our cryo-EM structures suggest that a tetramer-specific mechanism, a variation of the Venus flytrap model, can also engage. Specifically, in the intermediate-form A2Moo tetramer, TED within the induced protomer is shifted to interact with MG1 of the adjacent native-form protomer (Figure 3, Figure S6). This inter-protomeric MG1-TED interaction is also observed in the induced-form human A2M crystal structure (Marrero et al., 2012). This new lateral inter-protomeric interaction gained upon the cleavage-induced transformation may stabilize the induced-form A2M and prevent the prey chamber from re-opening (Figure 7B).

### Amine-induced A2M transformation

It has long been known that, not only bait region cleavage, but also small amines that inactivate thioester cause A2M tetramer transformation (Barrett et al., 1979). However, our model suggests that the A2M tetramer transformation induced by small amines may not be identical to the protease-induced A2M tetramer transformation. In the amine-induced A2M, the bait region should remain intact, blocking the structural transformation of MG2 and MG6 (Figure 5). This idea is also supported by the fact that the bacterial A2M monomers cannot be transformed by methylamine while it can capture proteases upon bait cleavage (Garcia-Ferrer et al., 2015; Wong and Dessen, 2014). This also suggests that amines can induce A2M transformation, specifically in tetrameric A2Ms. By reacting with thioester, small amines would weaken the hydrophobic interaction between RBD and TED, and release RBD from TED. Then, the inter-protomeric lateral MG1-TED interactions would be formed in the amine-induced A2M tetramer by bypassing the requirement of the cleavage-mediated shifting of MG2 and MG6 that detaches RBD from TED. Unfortunately, in the crystal structure of methylamine-induced human A2M, the bait region appeared to be cleaved, likely due to the month-long crystallization process (Marrero et al., 2012), preventing us from assessing this possibility from the structural comparisons.

### Number of proteases captured by A2M tetramer

Our intermediate-form A2Moo structure suggests that the structural induction of one out of four protomers within the tetramer is insufficient to trigger the global structural transformation (Figure 6A). Thus, one may assume that a single A2M tetramer can capture up to four protease molecules one by one. However, it was reported that A2M tetramer captures only two small protease molecules or a single large protease (Rehman et al., 2013). This conflict is not resolved by our cryo-EM structures. Since two grooves formed between chain A and chain D (or between chain B and chain C) are connected to form a large prey chamber, it is possible that a single protease molecule can cleave two baits in one prey chamber. However, failure to detect a tetrameric structure that possesses two induced protomers suggests that such processive bait cleavage by a single protease within the same chamber is rare.

### The structural difference between intermediate-form frog A2Moo and induced-form human A2M

The comparison between the induced-protomer of the intermediate-form frog A2Moo and the induced-form human A2M showed that the bulky finger module (MG7, CUB, TED, and RBD) in the induced-form human A2M is further shifted ~10Å toward the center of the tetramer (Figure 3C right). In the intermediate-form A2Moo, there is a larger space between the thumb module (MG3 and MG4) and the mobile bulky finger module (MG7, CUB, TED, and RBD) than in the induce-form human A2M (Figure 3C right). There are several hypotheses to explain this difference. First, the solved intermediate-form A2Moo may reflect a capture of a large protease in the *Xenopus* egg, whereas the crystal structure of human A2M purified from blood was induced by methylamine but not bound to a protease. Therefore, the bulky finger module may not be able to completely close the groove in A2Moo due to the presence of a protease. Second, the bulky finger module may not completely be closed until another or more protomer(s) in the tetramer is/are induced. Third, the induced-form human A2M structure may be artificially compacted due to the crystal packing. Fourth, the structural disparity may reflect amino acid sequence difference between frog A2Moo and human blood A2M (Figure S3). To solve this issue, the cryo-EM structure of a fully induced-form A2Moo tetramer structure would be needed.

### Limited structural changes in RBD upon induction

Induced-form A2M tetramers bind to several types of receptors (e.g., LRP1 and CD91) on the cell surface to cause receptor-mediated endocytosis, which is essential for the clearance of protease-captured A2M tetramer and other A2M functionaries, such as clearance of misfolded proteins (e.g., Aβ peptide and Parkinson’s disease-associated alpha-synuclein) and transportation of cytokines and growth factors (Cater et al., 2019; Garcia-Ferrer et al., 2017; Rehman et al., 2013). RBD of A2M C-terminus is responsible for the receptor binding (Van Leuven et al., 1986b, 1986a). The immunoelectron microscopic study revealed that the RBD is exposed to the tetramer surface by the A2M tetramer transformation induced by chymotrypsin (Delain et al., 1988). This exposure of the RBD in induced-form A2M is expected to trigger the receptor binding (Cater et al., 2019; Garcia-Ferrer et al., 2017; Rehman et al., 2013). However, in the native-form A2Moo structure (Figure S7), most RBD surfaces are already exposed to the solvent. The only region additionally exposed in the induced-protomer of the intermediate-form A2Moo is the hydrophobic pocket region that is used for protecting thioester in the native-form A2Moo (Figure S7). This suggests that the region around the hydrophobic pocket, which binds TED in native-form A2M, may contribute to the receptor binding in the induced-form A2M.

Fertilization is a unique developmental process in which a host egg cell acquires the exogenous sperm DNA. We and others have recently reported that the cytoplasmic DNA sensor cGAS in the innate immune system cannot be activated by the self-DNA by nucleosome-dependent inhibition (Boyer et al., 2020; Cao et al., 2020; Kujirai et al., 2020; Pathare et al., 2020; Zhao et al., 2020; Zierhut et al., 2019), but sperm DNA, which lacks nucleosomes, does not stimulate inflammation likely due to the lack of cGAS in eggs (Wühr et al., 2014). As several pathogens can infect frog oocytes and eggs (Fernández-Benéitez et al., 2008; Haislip et al., 2011; Holz et al., 2015; O’Connell et al., 2011), while sperm and eggs secrete proteases, such as acrosin and ovochymase, respectively (Lindsay et al., 1999, 1992; Mao and Yang, 2013), further studies are required to establish the physiological roles and regulation of egg cytoplasmic A2Moo and its receptor.

## Method

### Cryo-EM data processing

The pipeline of the cryo-EM processing is depicted in Fig. S2. The previously reported cryo-EM micrographs of interphase and metaphase nucleosome from *Xenopus* egg extract chromosomes (EMPIAR-10691, EMPIAR-10692, EMPIAR-10746, and EMPIAR-10747) (Arimura et al., 2021) were motion-corrected and dose weighted with a binning factor of 2 using MOTIONCOR2 (Zheng et al., 2017) with RELION3.0 (Scheres, 2012). Topaz v0.2.3 (Bepler et al., 2019) was trained using the particles assigned to the Alpha-2-Macroglobulin family protein structure by Template-, Reference- and Selection-Free (TRSF) cryo-EM pipeline (Arimura et al., 2021), and 345,116 particles were picked by Topaz v0.2.3 (Bepler et al., 2019). Based on the result of 2D classification by CryoSPARC v3.1, particles were split into two groups: ‘alpha-2-macroglobulin class’ that is similar to A2M structure and ‘non-alpha-2-macroglobulin class’ that are dissimilar to A2M structure. Using *ab-initio* reconstruction of CryoSPARC v3.1, one class of alpha-2-macroglobulin 3D map was generated from ‘alpha-2-macroglobulin class,’ and four classes of decoy 3D maps were generated from ‘non-alpha-2-macroglobulin class’. Using these 3D maps, 3D classification (decoy classification) was performed with all picked particles by heterogeneous reconstruction of CryoSPARC v3.1. To improve the picking accuracy by centering the particles, 2D classification was performed in CryoSPARC v3.2 (Re-center 2D class = off, classes = 50), and off-centered particles were removed, retaining 55,655 particles. Particles were aligned to center by homogeneous refinement in CryoSPARC v3.2 applying D2 symmetry using the 55,655 particles, and then re-extracted by recentering in CryoSPARC v3.2 (with aligned shifts = on). Extracted particles were used for retraining Topaz v0.2.4 (downsampling factor = 8, expected number of particles =30). 374,358 particles were picked by Topaz v0.2.4 (radius of Topaz extracted regions = 10, particle threshold = −2), and 68,957 ‘alpha-2-macroglobulin class’ particles were purified *in silico* by two rounds of decoy classification with eight decoy maps using CryoSPARC v3.2. To isolate the structural variants of A2M, six new 3D maps were generated by *ab-initio* reconstruction from the particles assigned to the alpha-2-macroglobulin 3D map, and further 3D classification was performed with these six maps and 68,957 ‘alpha-2-macroglobulin class’ particles. As a result, three “native-form” maps (Class 1, 2, and 4 shown in Figure S1), one “intermediate-form” (Class 3 shown in Figure S1) map, and two low-resolution classes were obtained.

For native-form A2M tetramer, 48,823 particles assigned to the class 1 and 2 were used for non-uniform refinement of CryoSPARC v3.2 (D2 symmetry). The resolution (FSC 0.143 cutoff) of native-form A2M tetramer was 4.1Å in the local refinement. A mask to cover the protomer of the native-form A2M was generated, and local refinement of CryoSPARC v3.2 (D2 symmetry) was performed for the symmetry expanded particles. The resolution (FSC 0.143 cutoff) of native-form A2M protomer was 3.7Å in the local refinement.

For intermediate-form A2M tetramer, 11,170 particles assigned to the class 3 were used for non-uniform refinement of CryoSPARC v3.2 (C1 symmetry). The resolution (FSC 0.143 cutoff) of intermediate-form A2M tetramer was 6.4Å in the non-uniform refinement.

For 3DVA of the native-form A2M tetramer, 52,445 particles assigned to the class 1, 2 and 3 were used for non-uniform refinement of CryoSPARC v3.2 (D2 symmetry). After symmetry expansion, 3DVA of CryoSPARC v3.2 (Modes = 4, Filter resolution = 10Å, output mode = simple) was performed.

For the 2D classification shown in Figure 2B and S4A, 48,823 particles assigned to the class 1 and 2 were used for 2D classification of CryoSPARC v3.2 (50 classes).

Local resolutions of maps were calculated with CryoSPARC v3.2 (Figure S3A). All maps were sharpened by the Auto-sharpen of Phenix (Afonine et al., 2018). 3D Fourier shell correlations were calculated by remote 3DFSC processing server (Zi Tan et al., 2017) using the sharpened maps (Figure S3A).

### Protein identification, atomic model building, and refinement

The tentative atomic model of monomer A2M was built using the amino acid sequence of *Xenopus laevis* ovostatin (XP_041426100.1), and side-chain EM densities were manually investigated to find the A2M homolog that fits the EM map (Figure S1). Using the Swiss model (Waterhouse et al., 2018), the atomic model of XP_041426100.1 was generated based on induced-form human A2M crystal structure (Marrero et al., 2012). Atomic coordinates of each domain were extracted and fitted to the EM density of the locally-refined native-form A2Moo protomer one by one manually using Chimera (Pettersen et al., 2004) and refined by Phenix and Coot (Afonine et al., 2018; Emsley and Cowtan, 2004). By investigating the low conserved insertions, bulky residues, and aromatic amino acids, XP_018079932.1 (A2Moo) matched best to the EM density map. Using the Swiss model, the amino acid sequence of the tentative atomic model was replaced with A2Moo (Waterhouse et al., 2018). The replaced model was refined using Phenix and Coot (Afonine et al., 2018; Emsley and Cowtan, 2004). Thioester was added by restricting the bond length of the Cys958 SG atom and Gln959 CD atom to 1.7 Å during the Phenix refinement. N-linked glycoses were added to the extra-densities observed around asparagine side chains. In the local refined map, densities around the region interacting with neighboring A2M molecule and bait region were ambiguous, and atomic coordinates of these regions were not placed. The refined monomer atomic coordinates were then multiplied to fit the native-form A2M tetramer map. The atomic coordinates around the region interacting with neighboring A2M molecules were built. Using the low-passed (11Å) map of native-form A2M tetramer, atomic coordinates around the bait region were built.

To build the atomic model of intermediate-form, chain A of the native-form A2M tetramer was removed, and atomic coordinates of each domain were extracted and fitted to the EM density of intermediate-form A2Moo tetramer one-by-one manually using Chimera (Pettersen et al., 2004). The built model was refined by Phenix and Coot (Afonine et al., 2018; Emsley and Cowtan, 2004). Chimera (Pettersen et al., 2004), ChimeraX (Goddard et al., 2018), and the PyMOL (Ver 2.0 Schrödinger, LLC.) were used for figure preparations.

## Data Availability

Atomic coordinates have been deposited in the Protein Data Bank under accession code PDB 7S62, 7S63, and 7S64. Cryo-EM density maps have been deposited in the EM Data Resource under accession code EMD 24847, 24848, and 24849.

## Acknowledgments

We are grateful to Mark Ebrahim, Johanna Sotiris, and Honkit Ng for their technical advice and assistance for the Cryo-EM. We thank Elizabeth Campbell, Seth Darst, Ruby Froom, Atsushi Ikai, Hideaki Konishi, Mirjana Lilic, Leena Sen, and Rochelle Shih for comments to the manuscript. This work was conducted with the help of the High-Performance Computing resource center and the Evelyn Gruss Lipper Cryo-Electron Microscopy Resource Center at the Rockefeller University. This work was supported by National Institutes of Health Grants R35GM132111 to H.F. and by the Japan Society for the Promotion of Science Overseas Research Fellowships to Y.A..

## Author Contributions

Y.A. designed and conducted all analyses. Y.A. and H.F. wrote the paper.

---

**Figure S1.**
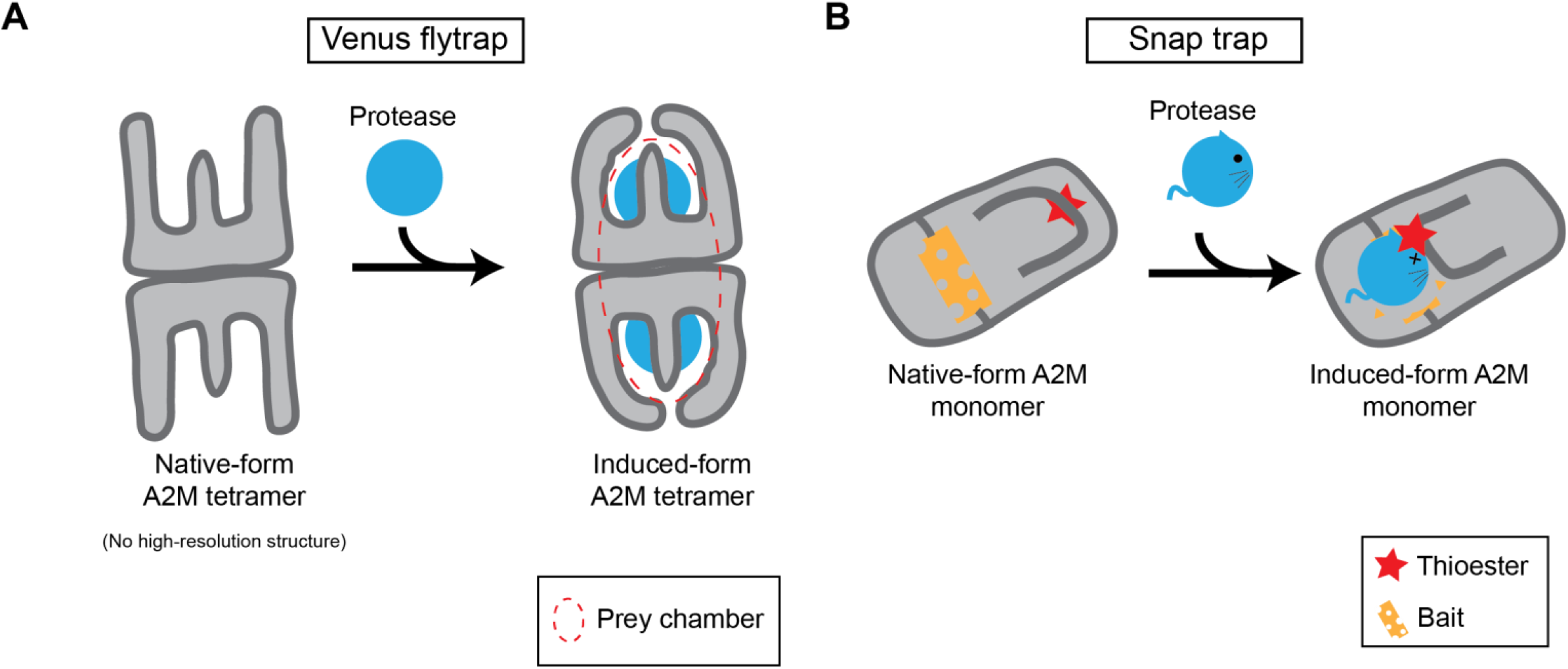
The Venus flytrap and snap trap mechanisms related to Introduction. (**A**) The Venus flytrap mechanism (**B**) Snap trap mechanism.

**Figure S2.**
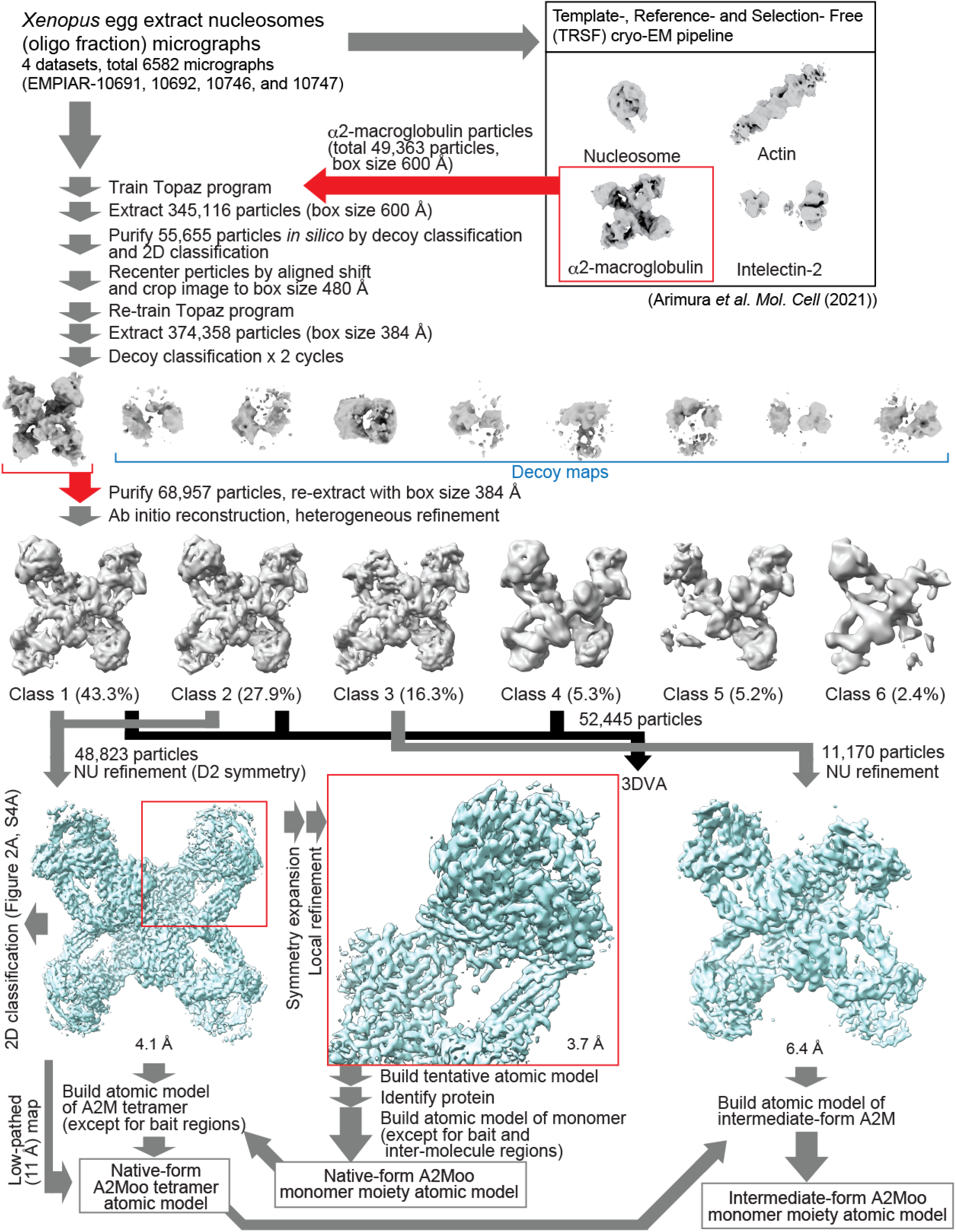
Cryo-EM analysis pipeline related to Figure 1. The pipeline of the cryo-EM data processing. See the method section for a detailed explanation.

**Figure S3.**
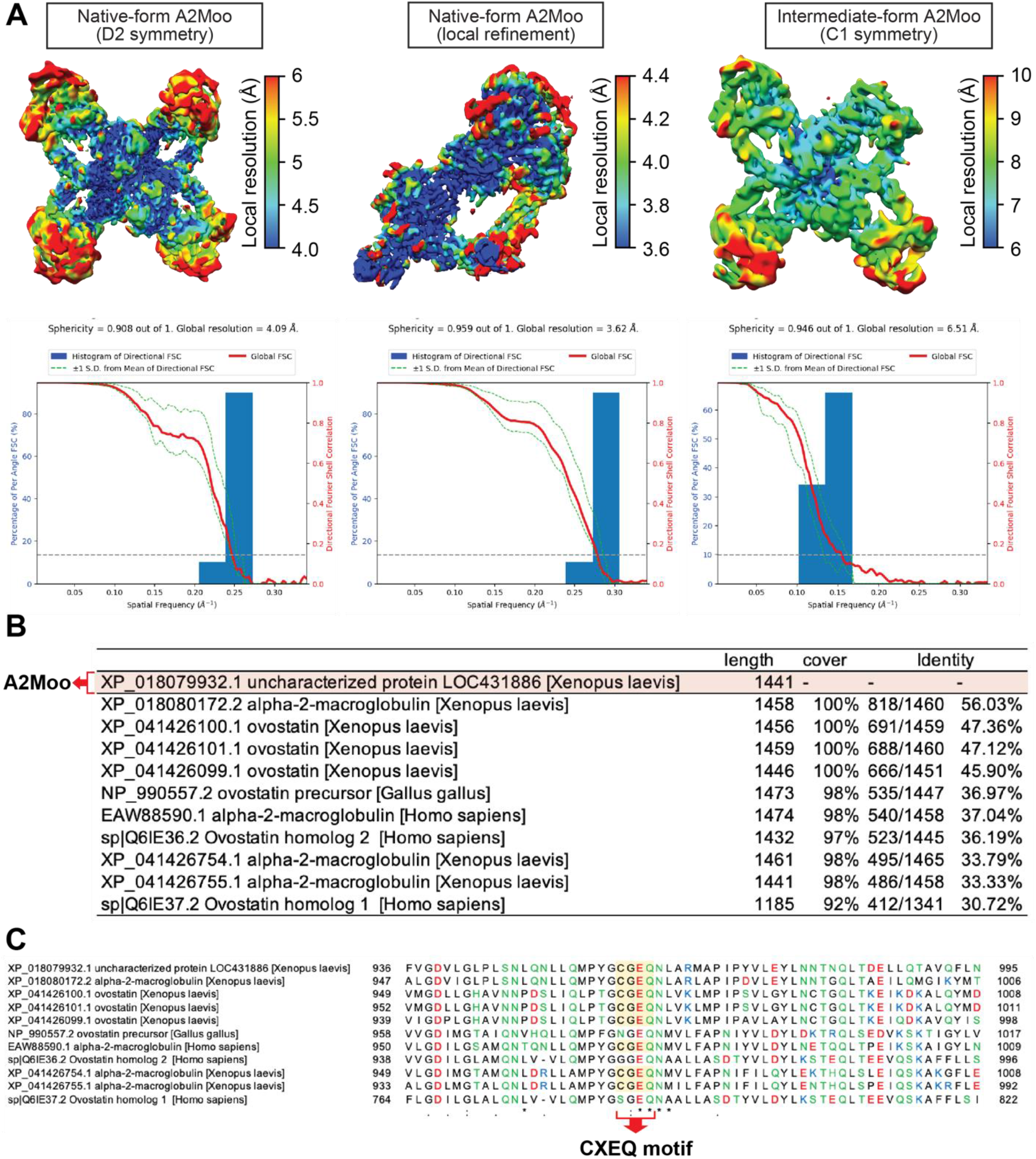
Cryo-EM structure determination of A2Moo related to Figure 1. (**A**) Local resolution maps and 3D Fourier shell correlations of the A2Moo cryo-EM structures. (**B**) Amino acid sequence conservation of A2Moo, A2M, and ovostatin encoded in human (*homo sapiens*), chicken (*gallus gallus*), and African clawed frog (*Xenopus laevis*). (**C**) Sequence alignment around CXEQ motif of A2Moo, A2M, and ovostatin encoded in human, chicken, and African clawed frog.

**Figure S4.**
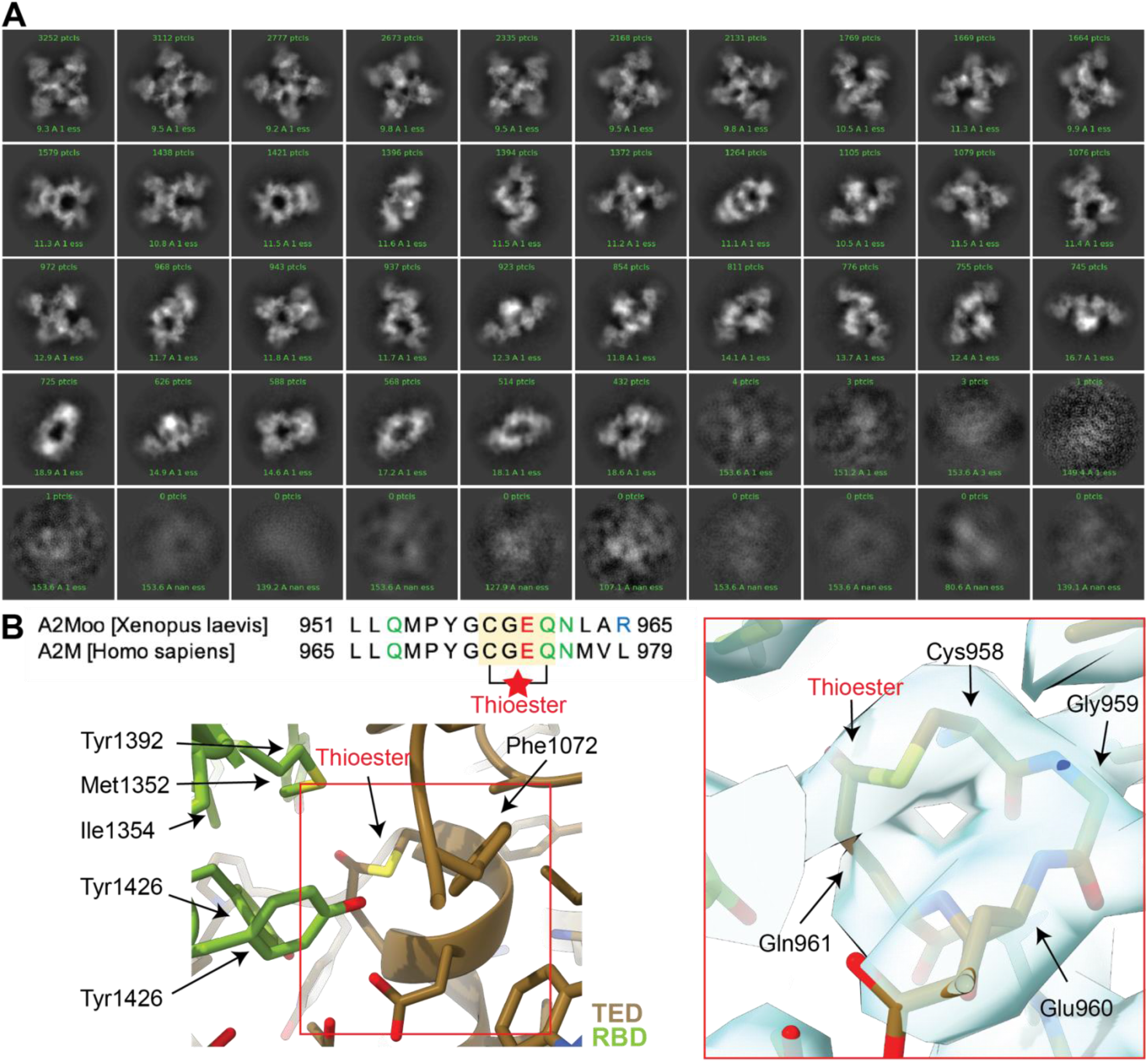
Cryo-EM structures of A2Moo tetramers related to Figure 2 and 3. (**A**) 2D class averages of the native-form A2Moo tetramer. All fifteen classes of the 2D classes of the native-form A2Moo tetramer are shown. (**B**) The cryo-EM density of around thioester in the native-from A2Moo. The locally refined maps of the native-from A2Moo and its atomic model are overlayed.

**Figure S5.**
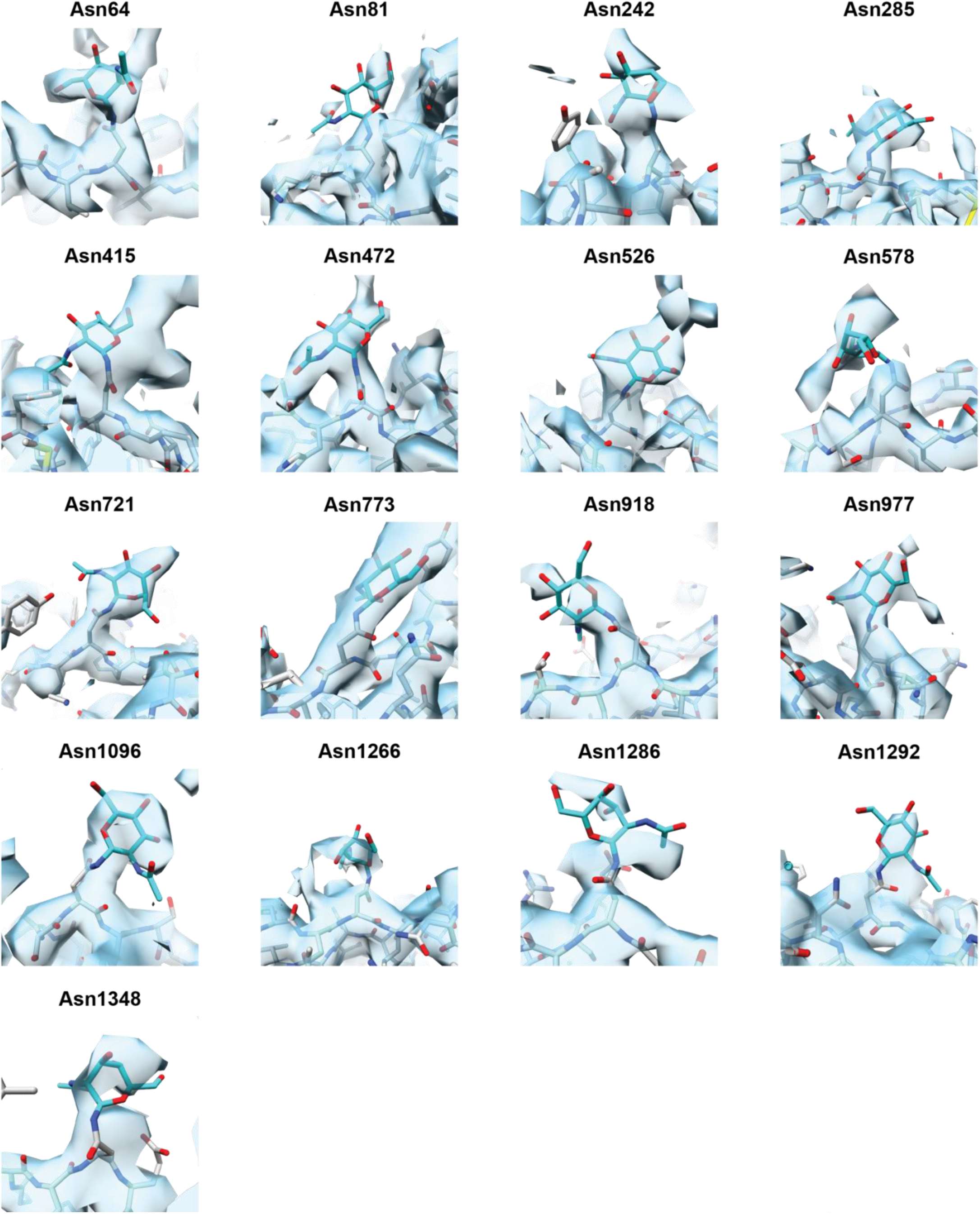
The cryo-EM density of around the *N*-linked glycosylation sites related to Figure 2 and 3. The locally refined map of the native-from A2Moo and the atomic model around the *N*-linked glycosylation sites are overlayed.

**Figure S6.**
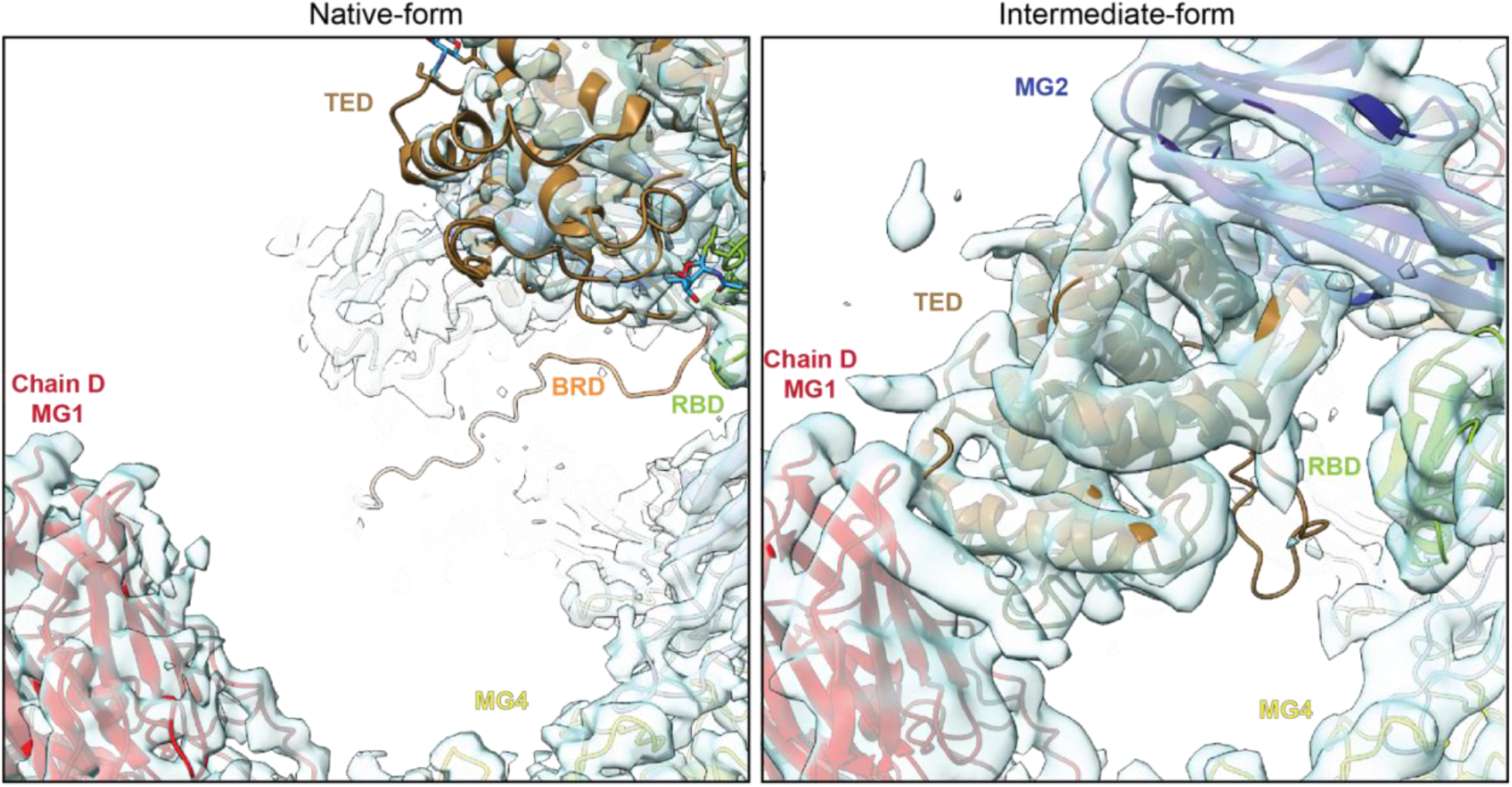
Structural comparison around lateral TED-MG1 interaction site related to Figure 2 and 3. Structural comparison around the lateral interprotomeric TED-MG1 interaction site between native-form (left) and intermediate-form (right) A2Moo.

**Figure S7.**
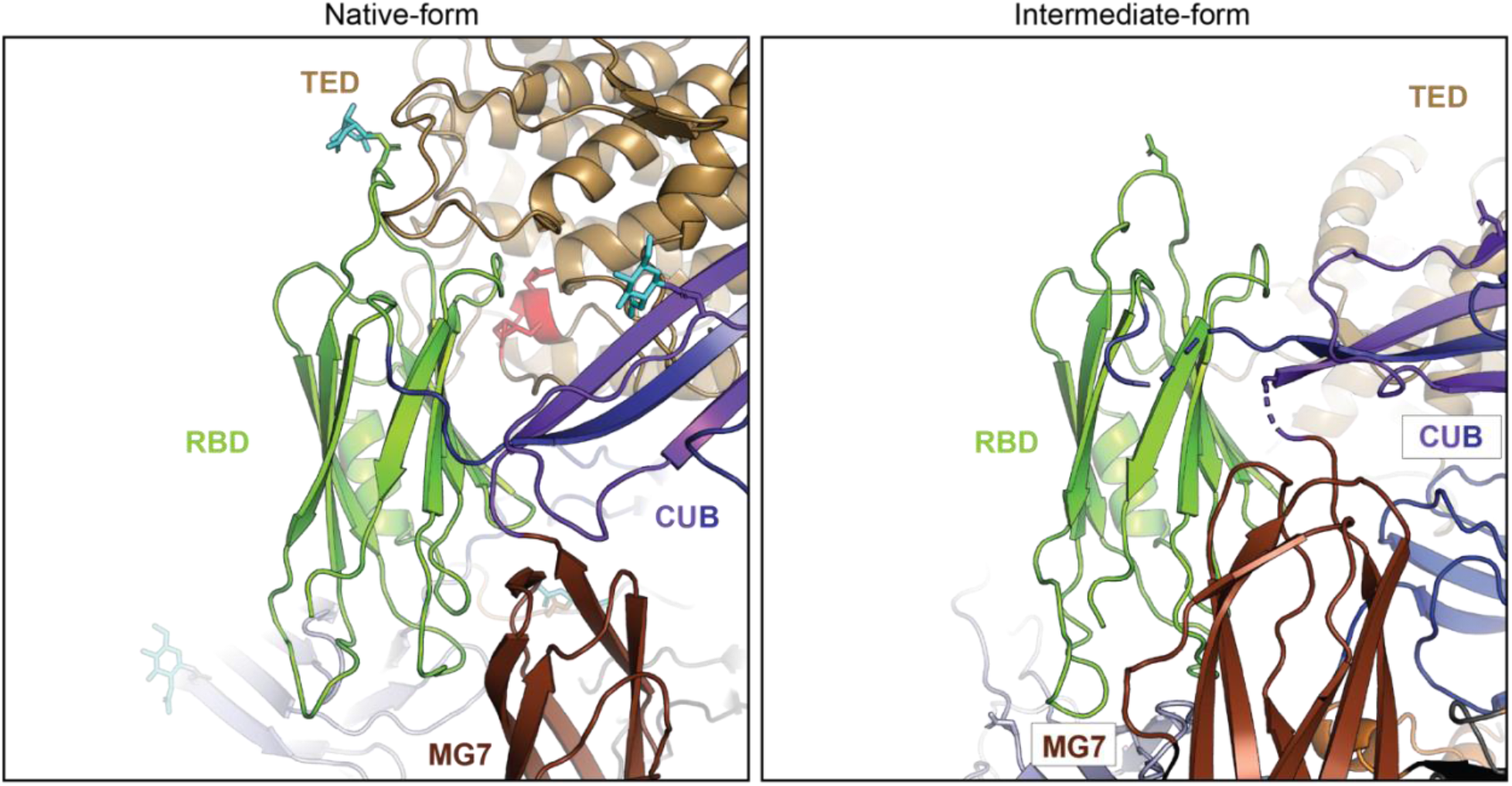
Structural comparison around RBG related to Figure 2 and 3. Structural comparison around RBG between native-form (left) and intermediate-form (right) A2Moo.

**Table S3.**
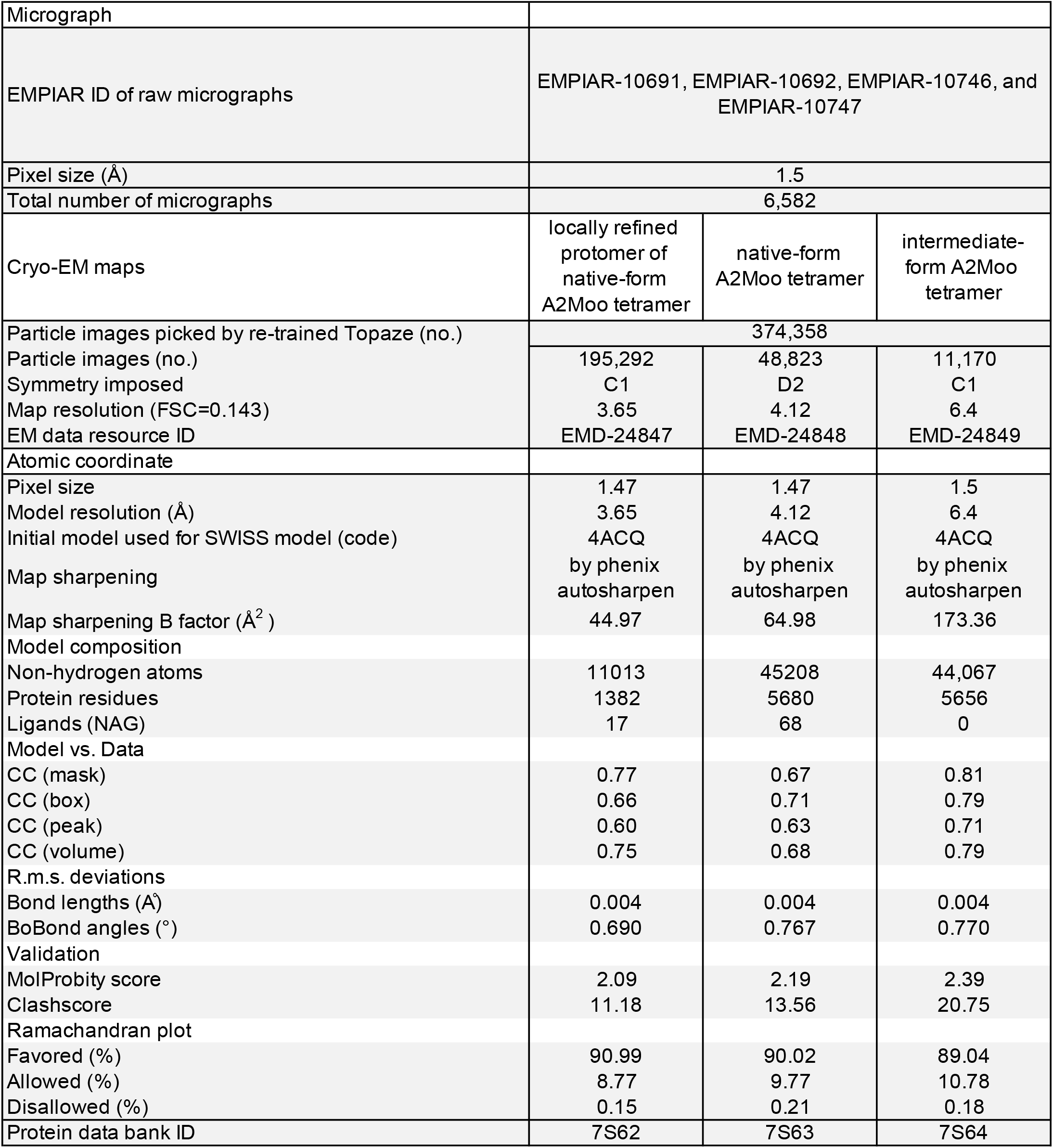
Data collection, model refinement, and validation

|  |  |  |  |
| --- | --- | --- | --- |
| Micrograph |  |  |  |
| EMPIAR ID of raw micrographs | EMPIAR-10691, EMPIAR-10692, EMPIAR-10746, and EMPIAR-10747 |  |  |
| Pixel size (Å) | 1.5 |  |  |
| Total number of micrographs | 6,582 |  |  |
| Cryo-EM maps | locally refined protomer of native-form A2Moo tetramer | native-form A2Moo tetramer | intermediate-form A2Moo tetramer |
| Particle images picked by re-trained Topaze (no.) | 374,358 |  |  |
| Particle images (no.) | 195,292 | 48,823 | 11,170 |
| Symmetry imposed | C1 | D2 | C1 |
| Map resolution (FSC=0.143) | 3.65 | 4.12 | 6.4 |
| EM data resource ID | EMD-24847 | EMD-24848 | EMD-24849 |
| Atomic coordinate |  |  |  |
| Pixel size | 1.47 | 1.47 | 1.5 |
| Model resolution (Å) | 3.65 | 4.12 | 6.4 |
| Initial model used for SWISS model (code) | 4ACQ | 4ACQ | 4ACQ |
| Map sharpening | by phenix autosharpen | by phenix autosharpen | by phenix autosharpen |
| Map sharpening B factor (Å<sup>2</sup>) | 44.97 | 64.98 | 173.36 |
| Model composition |  |  |  |
| Non-hydrogen atoms | 11013 | 45208 | 44,067 |
| Protein residues | 1382 | 5680 | 5656 |
| Ligands (NAG) | 17 | 68 | 0 |
| Model vs. Data |  |  |  |
| CC (mask) | 0.77 | 0.67 | 0.81 |
| CC (box) | 0.66 | 0.71 | 0.79 |
| CC (peak) | 0.60 | 0.63 | 0.71 |
| CC (volume) | 0.75 | 0.68 | 0.79 |
| R.m.s. deviations |  |  |  |
| Bond lengths (Å) | 0.004 | 0.004 | 0.004 |
| Bond angles (°) | 0.690 | 0.767 | 0.770 |
| Validation |  |  |  |
| MolProbity score | 2.09 | 2.19 | 2.39 |
| Clashscore | 11.18 | 13.56 | 20.75 |
| Ramachandran plot |  |  |  |
| Favored (%) | 90.99 | 90.02 | 89.04 |
| Allowed (%) | 8.77 | 9.77 | 10.78 |
| Disallowed (%) | 0.15 | 0.21 | 0.18 |
| Protein data bank ID | 7S62 | 7S63 | 7S64 |

